# The Impact Of Long-term Grazing Intensity On Functional Groups Richness, Biomass, And Species Diversity In an Inner Mongolian Steppe Grassland

**DOI:** 10.1101/2021.05.24.445414

**Authors:** Yousif Mohamed Zainelabdeen, Ahmed Ibrahim Ahmed, Ruirui Yan, Xiaoping Xin, Cao Juan, Jimoh Saheed Olaide

## Abstract

Livestock grazing is one of the major land uses, causing changes in the plant community's structure and grasslands composition. We assessed the effect of grazing intensity on aboveground biomass, species richness, and plant functional group (PFG) diversity in a temperature meadow steppe in Hulunbuir in northern China, involving 78 plant species from eight functional groups. Four grazing intensity classes were characterized, including light, moderate, heavy, and no grazing, based on stocking rates of 0.23, 0.46, 0.92, and 0.00 animal units per hectare. Our results show that the richness of short species, including perennial short grass, perennial short grass, and legume increased under light to moderate grazing, while no effect of grazing was observed on the richness of shrubs. With increasing grazing intensity, the aboveground biomass of perennial tall grasses and perennial tall forbs decreased significantly, while that of annual/biennial plant functional groups increased. The community diversity and evenness of annual/biennial plants increased significantly with grazing intensity. We concluded that heavy grazing has negative impacts on plant functional group richness and aboveground biomass.

## Introduction

Livestock grazing is considered one of the primary biotic factors that affect the natural grassland ecosystem functions (Olff and Ritchie 1998). Specifically, species diversity response to grazing intensity varies from location to location (De Bello, et al. 2006). This suggests that the magnitude of grazing intensity has the potential to impact species diversity, which often leads to vegetation degradation (Bakker 1998). Previous studies have shown that the positive impact of grazing intensity in natural grasslands implies that historical traditional grazing does not have a negative impact on these ecosystems (Montalvo, et al. 1993; Verdú, et al. 2007). For example, (Proulx and Mazumder 1998) reported that species richness responds positively to heavy grazing intensity in a highly productive environment. It is noteworthy, however, that grazing intensity is a complex disturbance that can affect the grassland communities, both directly and indirectly (Papanikolaou, et al. 2011), because it has the potential to change the vegetation structure, growth, and composition of different species, while also affecting the abiotic components of the ecosystem (Facelli and Springbett 2009). Whereas changes in grassland species composition and richness are induced by grazing intensity (Bergmeier, et al. 2003; Suding, et al. 2008), the influence of the latter on plant functional groups at the species level could differ (DÍAz, et al. 2007; Díaz, et al. 2001; McIntyre, et al. 1999; Noy-Meir, et al. 1989). However, these results were often inconsistent, more research is needed to improve our understanding of how grazing intensities affects species and plant communities (e.g., the species in the same plant functional group can respond differently to grazing intensity due to palatability or species structure), especially in Hulunbuir grasslands (Foggin 2008; Le, et al. 2014).

The disturbance caused by large herbivore grazing on plant functional groups’ life is an important factor affecting plant community composition. In some environments, the composition and structure of plant and animal species are influenced by grazing intensities, while in other places, the environmental characteristics of the area, including weather conditions and climate, are sometimes more important. Grazing directly influences plant community composition through the physical removal of plant parts. It can also indirectly influence plant communities by organizing ecosystem productivity or changing nutrient patterns between plants of different sizes. Therefore, grazing can change the quantity, diversity, and distribution of organisms in the ecosystem. Grazing intensities also influence grass species performance and grassland environment (Saccone, et al. 2014)

Biodiversity is a complex concept that allows for a variety of possible definitions. In a broad sense, from species to various forms of life, are defined as ecosystems (Hill 1973; Wilson, et al. 2016). At the community level, the diversity index reflects the distribution of the basic plant species in an environment. However, the term ‘diversity index’ could also be used to depict other categorizations such as functional groups and genetic types within the ecological environment. The entities of interest are usually individual animals or plants, where individual species, biomass, or coverage can be used as a measure of abundance (Dumont, et al. 2009; Fanselow, et al. 2011) It has been reported that grazing intensities had different impacts on single species’ aboveground biomass and biodiversity, which is documented by significant effects of grazing intensity on biomass and species diversity for this study.

Diversity can be divided into three parts: α diversity, which is the richness of a single site or specific community; β diversity, which is the difference of flora between different sites or communities; and γ diversity, which is the difference between geographical regions (Sepkoski Jr 1988). α diversity measures the aggregation within the community and reflects the fine division of ecological resources by species, while β diversity can be used to measure the degree of species composition turnover in the environmental gradient, potentially reflecting the degree of habitat selection or allotment. Gamma and beta diversities are similar, although gamma is measured at a larger spatial scale; it reflects the locality or degree of locality of a biota. (Whittaker and Rh 1977) retained the term “gamma diversity” for the combination of alpha and beta diversity, which he also called “complete diversity” (Whittaker 1972).

Palatability refers to the characteristic or condition of plant stimulating animal selective response, also defined as a pleasant taste. Preference can be expressed in the relative of time that animals graze different species. Preferences were left to animals to choose from, basically behavioral. Relative preference refers to the proportion choice between two or more species. In addition to palatability, many other factors affect food selection (Heady 1964).

Broadly, the objective of this study was to understand how different grazing intensities influence species richness, diversity and plant functional groups in the Hulunbuir meadow steppe ecosystem. Our goal was to determine the effect of grazing intensity on aboveground biomass and species richness based on species palatability. We hypothesized that perennial tall grasses are more palatable than other functional groups (e.g., shrubs are spiny, and Liliaceae are often poisonous). Plant functional groups are greatly used in community ecology and earth system simulation to describe the changes in plant characteristics within and between communities. However, this approach assumes that functional groups can account for a large proportion of variation in traits among species. Therefore, the use of functional groups is essentially a trait-based approach. Plant species in functional groups have similar characteristics and ecological similarity (Thomas, et al. 2019).

Furthermore, we evaluated the response of plant functional groups (PFGs) diversity to grazing intensity. Here, we hypothesize that the diversity of functional groups differs across grazing intensities. To clarify, species diversity implies a measure of both species number and abundance while species richness is just the number of plant species found in an environment.

## Materials And Methods

### Study area

The research was conducted in the Hulunbuir meadow steppe in northeast Inner Mongolia of China. The Hulunbuir meadow steppe has an area of 7.86 × 10^4^ square kilometers (Figure 1).

**Fig.1.**
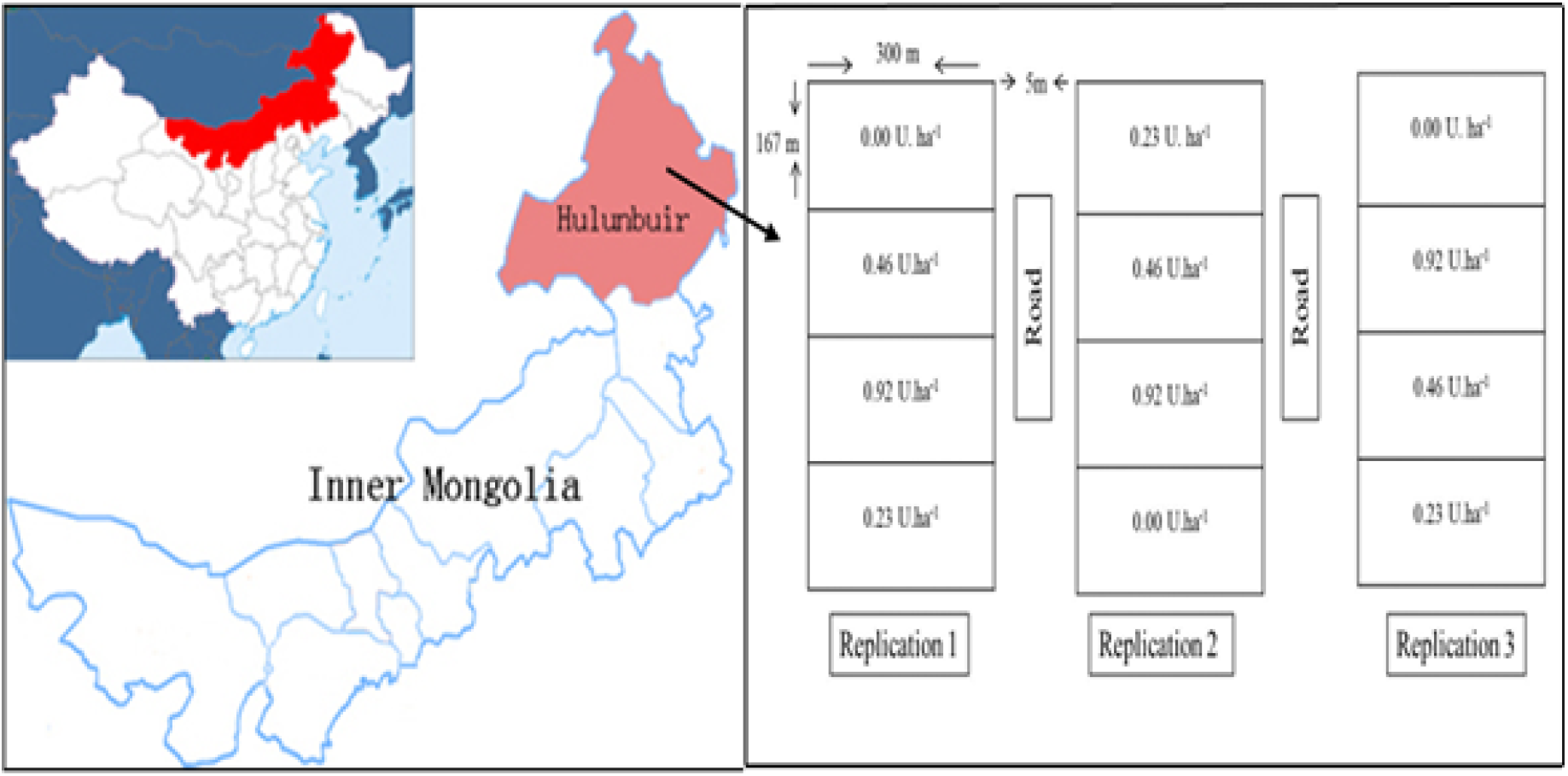
The geographical location of the Hulunbuir meadow steppe.

The mean maximum temperature in Hulunbuir is measured at 36.17 °C in July, and a minimum temperature of −48.5 °C in January. The annual frost-free period ranges from 85 to 155 days. Sunshine duration accounts for 2650-3000 hours per year. The area receives an average annual precipitation of 350 to 400 mm, 80% of which falls between July and September. Chernozem soils are commonly found here. The most dominant grass species in Hulunbuir are *Leymus chinensis, Carex duriuscula, Pulsatilla turczaninovii, Stipa baicalensis, Cleistogenes squarrosa, Artemisia tanacefolia* and *Thalictrum squarrosum*.

### Experimental Layout

The experiment was carried out on 600 hectares of natural grassland, with three grazing intensities and a no-grazing control (NG). The grazing intensities were 0, 0.23, 0.46, and 0.92 animal unit (AU) ha-1 for NG, light grazing (LG), moderate grazing (MG), and heavy grazing (HG), respectively. A 500 kg adult cattle is defined as 1 AU. The NG plot has been fenced and excluded from grazing since 2014. Each treatment plot occupies 5 ha of grassland with three replicates. In total, 12 plots were randomly distributed. The experiments were conducted during the grazing seasons (June to October) in 2014-2017. Drinking water was provided from an external source for the anmals. The numbers of animals grazed in each of the experimental plot was 0, 2, 4 and 8 heads of 250-300 kg calves for the treatment of NG, LG, MG and HG, respectively.

In each treatment plot, five sampling quadrats of 1 m2 were randomly located. Plants in each quadrat were trimmed to 2.5 cm in height. We divided the harvested species parts trimmed off into eight PFGs: perennial tall grass, perennial short grass, shrubs, legumes, Liliaceae herbs, annual/biennial plants, perennial tall forbs, and perennial short forbs (Ma, et al. 2010). Based on our sampling records, we estimated the species richness. To explore potential compositional changes related to grazing intensity, we also estimated the diversity index of each PFG, defined as the ratio of the rate of the abundance of species belonging to a particular functional group to the rate of occurrence of all the species captured in each quadrat.

The following formulas were used to calculate alpha diversity (Shannon–Wiener diversity index (H)), the Beta diversity index (β), Pielou evenness index (E), and important value (IV).

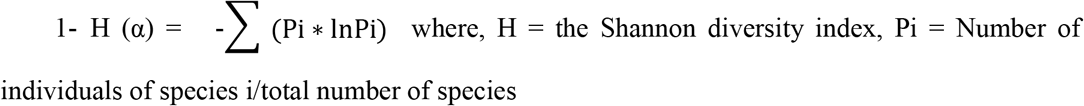

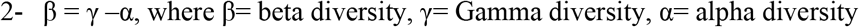

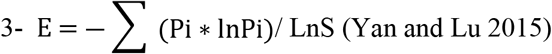

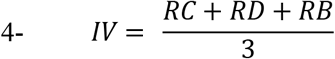

where RC is the relative coverage of the total species, RD is the relative density and RB is the relative biomass. Gamma diversity, also known as site diversity, is the total species richness of a site (Zhang, et al. 2014).

### Data analysis

We used a one-way analysis of variance (ANOVA) to examine the changes in species richness and diversity of each PFGs (all plant species) across the grazing intensity treatments using the SAS 9.2 (SAS, 2007). Duncan’s multiple comparison test was used to compute the significant difference among the treatments at *p* < 0.05. Similarly, for each replication, the difference in the composition of the dominant species and the 8 PFGs was tested by ANOVA.

## Results

### Grazing impact on species richness

The species richness of the PFGs as influenced by grazing intensity were summarized in Table 1. A significant difference (*p* <0.05) was observed between the treatments for perennial short grass (PSG), perennial tall forbs (PTF) and Liliaceae herb (LILY). Perennial short grass species were higher in the LG and MG plots (Table 1). With an increase in grazing intensity, species richness of PSG decreased under HG but recorded a significantly similar value with the NG treatment. The species richness of PTFs and LILY was higher at sampling plots subjected to light and moderate grazing, as well as the NG treatment. Further, heavy grazing reduced the richness of these species (Table 1).

**Table 1.**
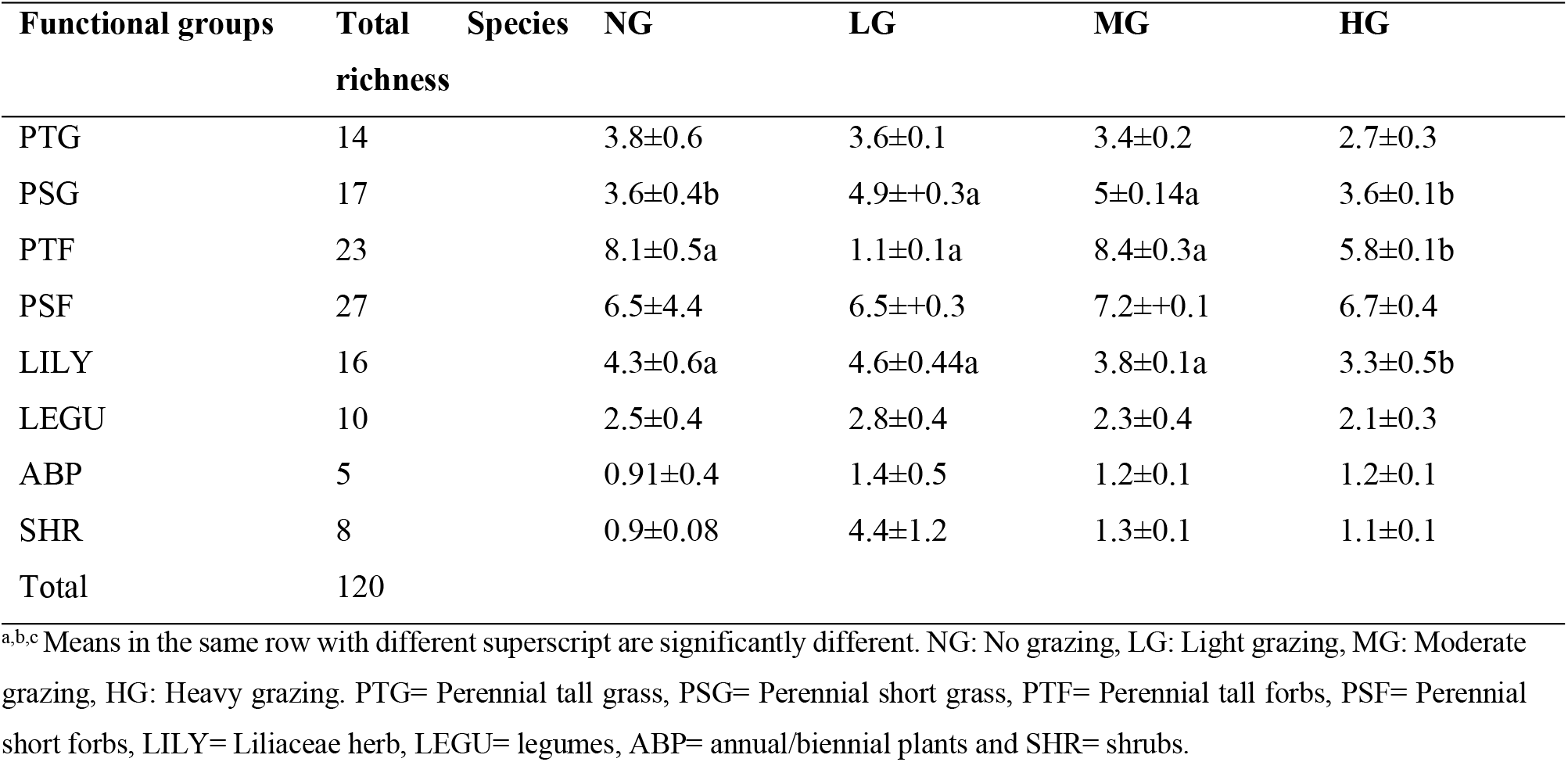
Species richness of plant functional groups under four grazing intensities.

### Response of plant functional groups’ aboveground biomass to grazing intensity

The aboveground biomass differed significantly (*p* <0.05) among the grazing intensities (Table 2). The NG plot had the highest PTG (102.0 g/m2) biomass while the least was recorded for MG (21.0 g/m2) and HG (4.0 g/m2) plots. Similarly, PTF (47.4) and LILY herbs (14.7) biomasses were higher in the NG plot and the lowest was recorded in the HG (10.9 and 0.98 g/m^2^) plot. The ABP functional group strongly increased in the HG intensity plot (2.2) while the NG (0.7) and MG (0.43) intensity plots recorded similar lower values. All the grazing intensity plots had higher SHR biomass than the NG plot. However, there were no significant differences (*p* >0.05) in perennial short forbs (PSG), PSF, and LEGU biomass across the grazing treatments imposed.

**Table 2.** Responses of aboveground biomass g/m^2^ of different plant functional groups to gradients of grazing intensity. Mean values and (± SE) for each group of plots for different plant functional groups

| Functional groups | Species number | NG | LG | MG | HG |
| --- | --- | --- | --- | --- | --- |
| PTG | 6 | 102.0±12.9a | 54.6±6.1b | 21.0±2.9c | 4.0±1.1c |
| PSG | 7 | 26.3±3.8 | 33.3±4.7 | 36.4±1.1 | 24.6±4.7 |
| PTF | 18 | 47.4±3.9a | 40.8±2.6ab | 30.3±5.2b | 10.96±1.5c |
| PSF | 16 | 17.9±4.4 | 13.5±1.1 | 14.6±2.9 | 11.2±0.95 |
| LILY | 8 | 14.7±5.3a | 11.4±0.7ab | 4.0±0.6bc | 0.98±0.1c |
| LEGU | 9 | 3.6±1.5 | 3.6±2.6 | 3.3±2.1 | 1.5±0.2 |
| ABP | 11 | 0.7±0.4b | 0.98±0.4ab | 0.43±0.07b | 2.2±0.57a |
| SHR | 3 | 1.2±0.4b | 4.4±1.2a | 5.6±0.09a | 4.8±1.3a |
| Total | 78 |  |  |  |  |

<sup>a,b,c</sup> Means in the same row with different superscript are significantly different. NG: No grazing, LG: Light grazing, MG: Moderate grazing, HG: Heavy grazing. PTG= Perennial tall grass, PSG= Perennial short grass, PTF= Perennial tall forbs, PSF= Perennial short forbs, LILY= Liliaceae herb, LEGU= legumes, ABP= annual/biennial plants and SHR= shrubs.

With grazing intensity, the decrease of dominance species was higher in PTG and PTF (3.85%), while the PSG and PSF dominant species increased with grazing intensity 3.85% and 8.97% respectively (Table3). Most of the C4 photosynthetic pathway dominant species appeared in PTF (5.13%) and PSF (2.56%). About 17.94% of the total grassland species showed a positive reaction to grazing intensity by increasing their aboveground biomass (Table 3), while 16.66% showed a negative response to grazing intensity by decreasing their aboveground biomass.

**Table 3.**
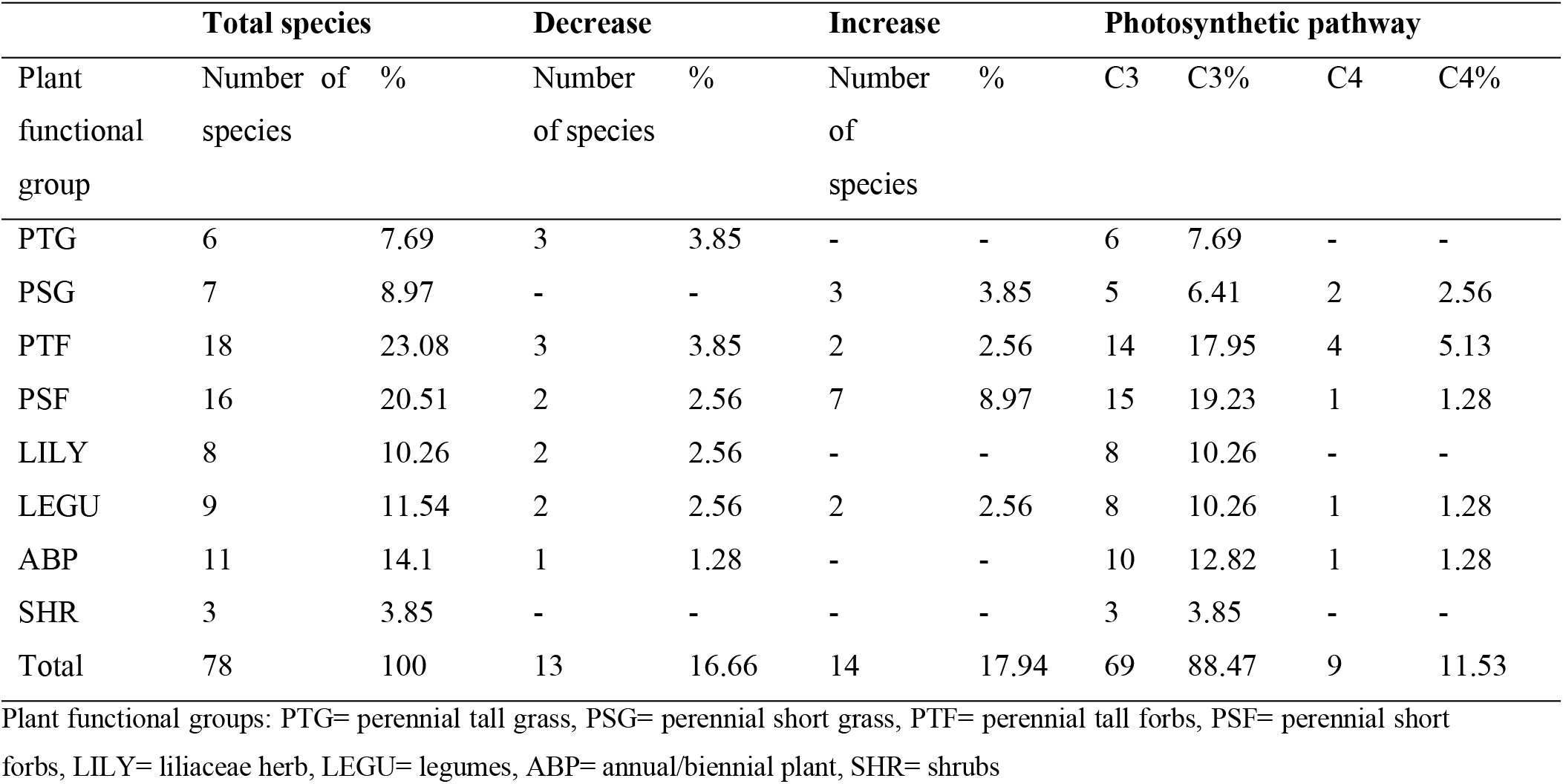
Proportional increase and decrease in the aboveground biomass and species photosynthetic pathway of plant functional groups across different grazing intensities

In this study, a total of 78 species were identified including 13 dominant species (Table 4). These species generally decreased with increasing grazing intensity. Specifically, we identified four palatable species (*Leymus chinensis, Bromus inermis, Achnatherum sibiricum,* and *Vicia amoena*) and three of them belong to the PTG functional group. There were six species with medium palatability (*Astragalus melilotoides, Lilium tenuifolium, Veronica i*ncan*a, Galium verum, Clematis hexapetla,* and *Thalictrum petaloideums*) belonging to LEGU, LILY herbs, ABP, and PTF functional groups respectively.

**Table 4.**
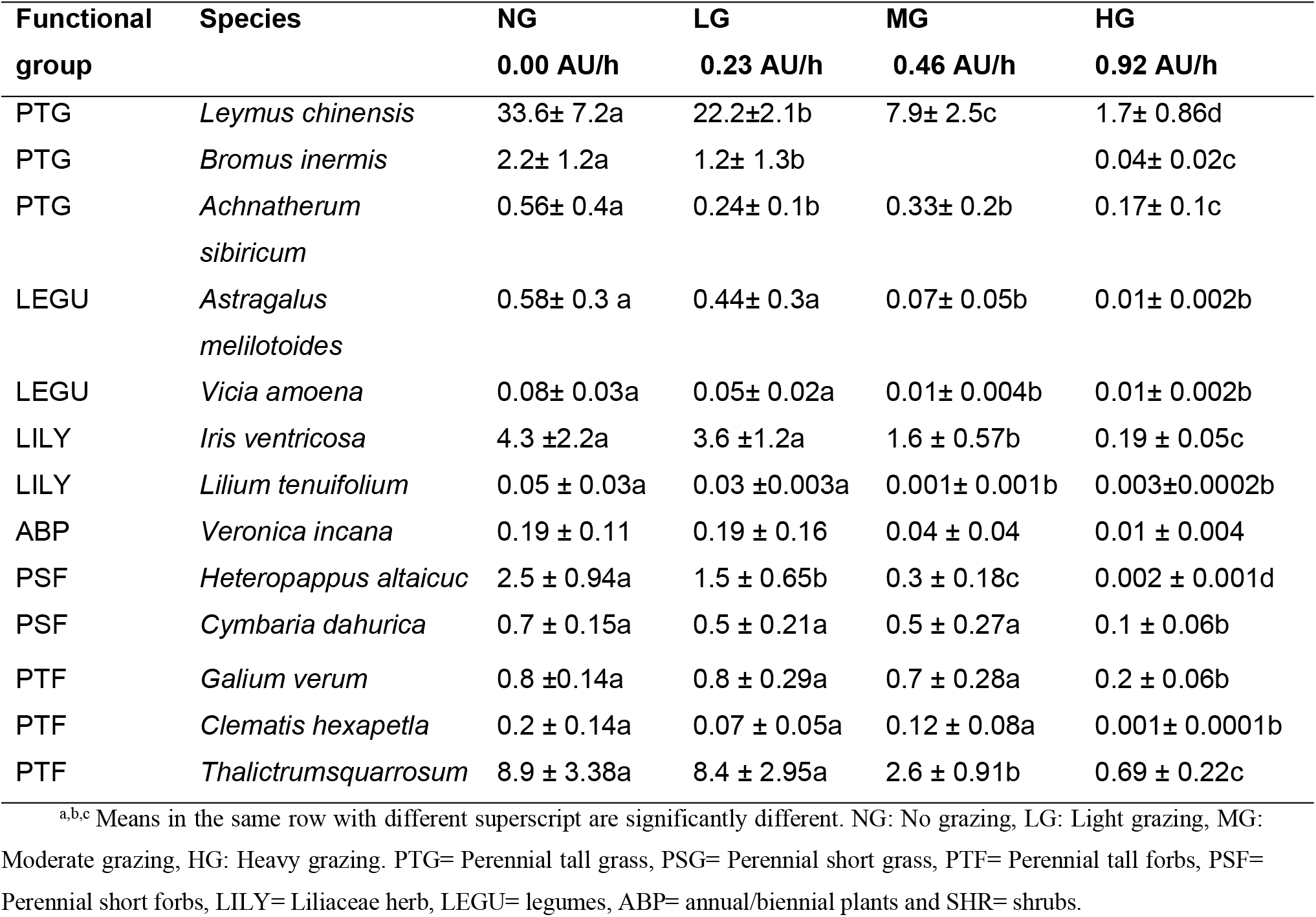
Variation of dominant species with high palatability in different plant functional groups across different grazing intensities

Table 5 shows plant functional groups with low to medium palatability. Generally, these plant species increase with an increase in grazing intensity.

**Table 5.** Variation of dominant species with low and medium palatability in different plant functional groups across different grazing intensities

| Functional group | Species | NG<br>0.00 AU/h | LG<br>0.23 AU/h | MG<br>0.46 AU/h | HG<br>0.92 AU/h |
| --- | --- | --- | --- | --- | --- |
| PSG | *Carex duriuscula* | 12.4 ± 3.5d | 20.8 ± 3.1c | 29.6 ± 4.05b | 42.8 ± 3.5a |
| PSG | *Cleistogenes squarrosa* | 1.3 ± 0.66 c | 3.5 ± 0.76b | 4.4 ± 1.1b | 6.2 ± 1.7a |
| PSG | *Koeleria cristata* | 0.61± 0.52b | 0.63 ± 0.20b | 1.8 ± 0.70a | 2.02 ± 0.61a |
| LEGU | *Astragalus adsurgens* | 0.04 ± 0.033c | 0.33 ± 0.15b | 0.58 ± 0.17a | 0.98 ± 0.25a |
| LEGU | *Oxytropis myriophylla* | 0.04 ± 0.04c | 0.02 ± 0.001c | 0.13 ± 0.09b | 1.0 ± 0.44a |
| PSF | *Pulsatilla turczaninovii* | 4.3 ± 0.97b | 4.3 ± 0.81b | 7.6 ± 1.2a | 7.1±1.3a |
| PSF | *Potentilla acaulis* | 0.07 ± 0.04c | 0.73 ± 0.41b | 1.00 ± 0.6b | 3.03 ± 1.1a |
| PSF | *Potentilla verticillaris* | 0.11± 0.01b | 0.16± 0.07b | 0.33± 0.16b | 0.71± 0.13a |
| PSF | *Taraxacum mongolicum* | 0.001±0.0001c | 0.09 ± 0.010b | 0.09 ± 0.05b | 1.4 ± 0.5a |
| PSF | *Potentilla tanacetifolia* | 0.03 ± 0.02b | 0.09 ± 0.03b | 0.30 ± 0.20a | 0.70 ± 0.32a |
| PSF | *Sibbaldia adpressa* |  | 0.04 ± 0.02b | 0.09 ± 0.07b | 1.02 ± 0.23a |
| PTF | *Artemisia tanacetifolia* | 9.1± 1.47a | 10.9 ± 2.48a | 11.7± 2.90a | 11.3 ± 2.17a |
| PTF | *Schizonepeta multifida* | 0.17±0.12 b | 0.22± 0.07b | 1.3± 0.5a | 0.97±0.43 a |

<sup>a,b,c</sup> Means in the same row with different superscript are significantly different. NG: No grazing, LG: Light grazing, MG: Moderate grazing, HG: Heavy grazing. PTG= Perennial tall grass, PSG= Perennial short grass, PTF= Perennial tall forbs, PSF= Perennial short forbs, LILY= Liliaceae herb, LEGU= legumes, ABP= annual/biennial plants and SHR= shrubs.

### Response of functional groups diversity to grazing intensity

Gamma species diversity and its composition (Alpha and Beta) were consistently higher in PTG, LEGU, ABP, SHR, and lower in PSG and PSF (Fig. 2). The contribution of β diversity to gamma diversity in Hulunbuir grassland is slightly greater than that of α diversity (Fig. 2b and c). The contribution of α and β diversity of the eight PFGs across the grazing intensities to γ diversity was different (Fig. 2a-2c). Species diversity was higher in PTF and PSF than other functional groups. The high diversity in PTF and PSF was mainly due to the higher number of species richness in these functional groups. We also found a significant difference (*p* < 0.05) in Pielou evenness of the eight PFGs across the grazing intensities (Fig. 2d). The evenness index was generally higher for PSF and LILY herbs across the grazing intensities, while PTG and LEGU recorded the lowest values in the HG treatment.

**Fig.2.**
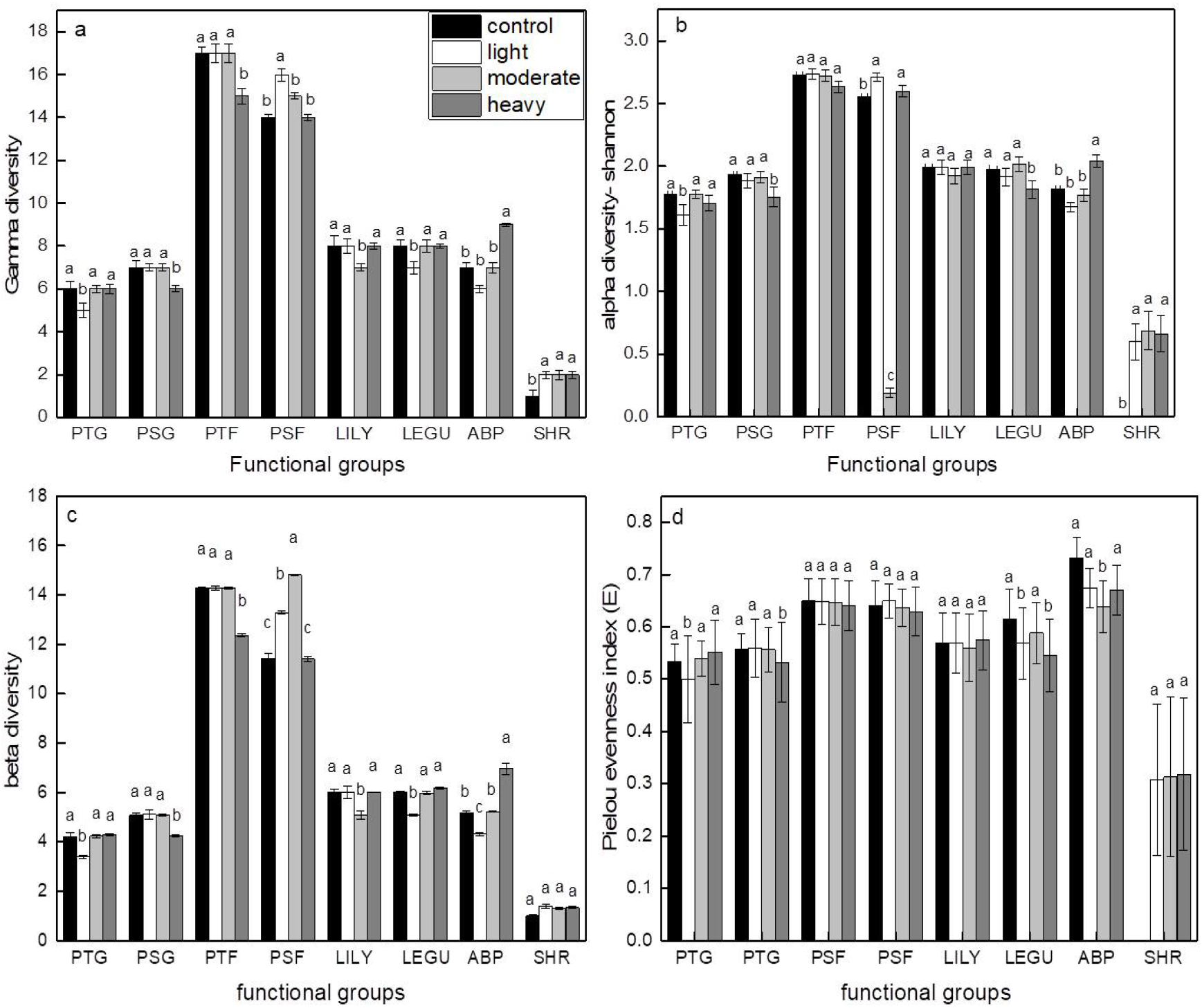
Comparisons of (a) Gamma diversity (γ), (b) Alpha diversity (α), (c) Beta diversity (β), and (d) Pielou Evenness of eight plant functional groups across different grazing intensities in Hulunbur grassland, Inner Mongolia. PTG: Perennial tall grass, PSG: Perennial short grass, PTF: Perennial tall forbs, PSF: Perennial short forbs, LILY: Liliaceae herb, LEGU: Legume, ABP: Annual/biennial plant, SHR: Shrub. Error bars indicate the standard error ((± SE) of the mean (p ≤ 0.05).

## Discussion

Livestock grazing may modify plant community structure, thereby leading to changes in vegetation characteristics. Across the length of the grazing season, several species could grow and become established within the grazing environment. Our findings demonstrate that species richness was consistently higher under LG and MG compared with the NG and HG intensities. This may be related to grazing history and the rate of species disappearance during defoliation in the HG plot (Marty 2005; Noy-Meir, et al. 1989). The trampling and nibbling effect of cattle led to reduced biomass through the removal of tall species and consequently result in changes in competitive interactions, which enabled less aggressive species to grow and hence a higher species richness in the community (Sternberg, et al. 2000). Several studies have proposed that grazing intensity and biodiversity converge in a general ‘dominance disturbance theory’ (Grime 1979; Sternberg, et al. 2000). Our results on the relationship between grazing intensity and species richness support this theory. In sum, species richness increased in both LG and MG plots.

We found that the effect of LG and MG on species richness did not differ and this may be related to the presence of grazing tolerant species in the treatments. Species richness was observed to be lower under HG intensity. There was no visible effect of light grazing during the plant growth period. Transition in PFGs as a result of grazing intensity has been documented in the literature (Papanikolaou, et al. 2011; Wang, et al. 2019; Wu, et al. 2014b). Our analysis of functional groups showed a decline in the PTF in the HG plot only. Similarly, PSGs declined in the NG and HG plots. These results contradicted the earlier report by (Papanikolaou, et al. 2011) who reported that the richness of both perennial grasses and forbs increased with grazing intensity. The difference between the former and present study may be related to vegetation type (Wu, et al. 2014b). However, our result is similar to the findings by (Fernández Alés, et al. 1993). In contrast, shrubs and legumes functional groups were less affected by grazing intensity. Their persistence is related to the development of chemical and/or physical defense substances such as chemical secretions and capillaries, as well as the possession of spiny leaves. They are also less dependent on seed production than other species. This strategy allows for rapid growth and early establishment after the first rainfall and improves grazing tolerance.

Our results showed that the total aboveground biomass of the PFGs decreased with increasing grazing intensity, and this is consistent with earlier reports (Wang, et al. 2016; Wu, et al. 2014a). Moreover, the plant composition changed due to the differential response of the PFGs to grazing intensity, which is weaker at the community level than for individual species based on aboveground biomass. The change in plant community composition is broadly driven by the decline in the biomass of the palatable species (especially PTG functional groups) and the increase in the aboveground biomass of ABP plants. These results agree with the previous study by (Sternberg, et al. 2015) that an increase in grazing intensity increases the dominant species of Gramineae which comprises of the most palatable species of grasses and forbs. Notably, some functional groups increased (e.g., ABP and SHR) while others decreased (e.g., PTG, PTF, and LILY) as grazing intensifies. This may be attributed to the higher probability of the tall grasses being defoliated by the grazing animals, thereby increasing light interception by the understory plants (Pekin, et al. 2014). Also, the reduction of some functional groups could have increased the competitive ability of others. In grassland ecosystems, however, dominant species have been reported to exhibit significant resistance or tolerance to grazing through a series of adaptive mechanisms (e.g., secretion of chemical defense substances) when the grazing condition changes (Aboling, et al. 2008; Ruppert, et al. 2015).

Species with different photosynthetic pathways have a different physiological and ecological response to grazing (Wang 2002). Animals accidentally eat some plants while they graze. Animals eager for fresh grass may accidentally bite off the crown of toxic species when they have nothing else to eat. This may explain the decrease of LILY functional group with grazing intensities

Grassland management decisions must, therefore, take seasonal development patterns and grassland succession into account, both of which are closely related to photosynthetic pathways. In Hulunbuir grassland, grazing induces the abundance of C4 species at the early succession stage. This is because the C4 species has the potential to produce a sufficient quantity of seeds that match their high seed dispersal capacity.

In this study, we found only nine species with the C4 photosynthetic pathway, while the majority of others (67 in total) have C3 pathways. The relatively low quantity of C4 species could be premised on the relative abundance of species within the eight PFGs. Our results suggested that C4 species are common in forbs than other PFGs and this corroborates the earlier report by (Wang 2002) that grazing intensity significantly influenced the abundance of C3 plants in a 4-year trial. Whereas C3 plants decreased with increasing grazing intensity, we found that the C4 plant increased as grazing intensifies. Simultaneously, summer temperatures also increased, providing an alternative explanation for the observed increase in C4 plants.

In Hulunbuir grassland, the structure and composition of the species change over time and space. Our results indicated that the species composition and abundance of all the eight PFGs are dynamic, thus, the different PFGs provide fodders of different quality at the same time of grazing season. (Tserendash and Erdenebaatar 1993) describes the early summer flowering contents of the species *Koeleria cristata, Poa partenis,* and *Agropyron cristatum* in steppe ecosystems. Notably, at the same time of the year, *Carex spp* showed late flowering. More importantly, in the growing season, the feeding C4 plants showed compensatory growth after grazing.

The patterns of species grown under different water ecotype can help us to understand the community structure, diversity and productivity of temperate grassland (Fay, et al. 2002; Fowler 1986; Köchy and Wilson 2000; Tsialtas, et al. 2001; Weltzin and McPherson 1997). The dominant species, *Leymus chinensis, Bromus inermis, Achnatherum sibiricum* and *Vicia amoena* were relatively higher in grasses and legumes. In total, 78 plant species belonging to eight functional groups (16.66%) decreased, while 17.94% increased with grazing intensity. Livestock grazing plays a significant role in plant species diversity and vegetation composition of grasslands (Cingolani, et al. 2003; Pucheta, et al. 2004). Light and moderate grazing intensity recorded higher plant species diversity compared with the NG and HG intensity. This implies that HG intensity resulted in a decrease in the species diversity of Hulunbuir grassland (Kikoti and Mligo 2015). Further, our results showed a decrease and increase in the palatable (e.g., *Bromus inermis*) and less palatable (e.g., *Pulsatilla turczaninovii*) species as grazing intensified. This finding concurs with the report by (Ji, et al. 2020) who reported an increase in the unpalatable aboveground biomass in total aboveground biomass with increasing grazing intensity. Also, the observed trend can be attributed to the selective defoliation of palatable species by the grazing animals (Liu, et al. 2015; Venter, et al. 2019) which leads to a decline in competition for resources (Ma, et al. 2019) and this was more evident in the HG plot.

The intensity of grazing is a key determinant of the spatio-temporal pattern of grassland resource utilization by livestock. The impact of grazing on plant species diversity can also be explained relative to individual species’ responses because of the variation in plants resistance and tolerance to grazing. Specifically, *Leymus chinensis* and *Achnatherum sibiricum* which were palatable species accounted for the highest important value of biomass in the NG plot. With increasing grazing density, *Leymus chinensis* gradually decreased due to preferential selection by the grazing animals (Liu, et al. 2016), hence, it represents the lowest value of aboveground biomass among the dominant species in the HG treatment. As grazing intensifies, *Carex duriuscula* and *pulsatilla turczaninovii* gradually become the dominant species in the HG plot. Overall, it can be inferred that the response of the aboveground biomass of both the palatable and less palatable species differ across the grazing intensities adopted in this study.

Plant species with different growth and reproductive phenology may have different responses to grazing intensities across the length of the grazing season. In this study, the response of individual species to different grazing intensities is different; however, the same number of positive and negative reactions among different species did not affect plant species richness and diversity. In general, the response of vegetation to grazing intensity may be strongly dependent on or overridden by vegetation type (Wu, et al. 2014a). Therefore, grazing intensity is the most important grazing management variable affecting the plant community structure in grassland ecosystems (Hickman, et al. 2004).

The contribution of α and β diversity to γ diversity is the basis of understanding the composition of biodiversity (Jost 2007; Zhang, et al. 2014). There are different opinions on the relative importance of α and β diversity as a function of γ diversity. Whereas some researchers hold the view that α diversity is more important, others attached a higher value to β diversity while some groups of scholars believe that the two (i.e., α and β) are of equal importance (Jost 2007; Meynard, et al. 2011). In this study, we found that β diversity contributes to gamma diversity more than α diversity for all PFGs across the grazing intensity and that total diversity is greater in species-rich PFGs (e.g., PTG and PSF) than species-poor PFGs (e.g., SHR and PTF). The relative contribution of α and β to γ diversity of the grassland community depends on the extent of species diffusion and ecological heterogeneity (Chiarucci, et al. 2010; Crist and Veech 2006). Alpha diversity is more important in the community with a homogeneous environment and species with high diffusion potential, while β diversity is more important in the community with a heterogeneous environment and species with weak diffusion potential. The observed increased in diversity of the functional groups may be attributed to species richness and this strengthens the high diversity recorded in species-rich functional groups. The results of the functional group diversity recorded across the grazing intensities were consistent with those reported earlier (Carlson 2011; Harrison, et al. 2003; Schultz, et al. 2011).The differences in the evenness of the PFGs in the various grazing intensity treatments depends on the difference between the PFGs (Stirling and Wilsey 2001). Because species evenness index and richness might respond differently to different grazing intensity and environmental variables, we found different responses of plant functional groups to evenness. The grazing density had differential impact on diversity, even though 78 plant species were included in every season.

## Conclusions

In this study, various sensitive indices of long-term grazing were set and evaluated. Grazing intensity had a different effect on (change vegetation composition) depends on the productivity and plant functional groups of grassland communities. There was a significant negative effect of grazing intensity on the aboveground biomass (AGB). Therefore, the grazing intensity levels are inverted the grazing pressure from non-grazing to heavy grazing intensity. AGB can be used as a potential sensitive index for meadow productivity. In this study, LG and HG represent the best and worst responses of aboveground biomass to grazing, respectively. We found that the richness of the PFGs only differed for PSG and LILY across the grazing intensities. Perennial short grass species were higher in the LG and MG plots. The PTG showed significantly decreased with grazing intensity, while PSG increased with grazing intensity. Also, this study showed that the effect of grazing density on grassland diversity was consistently higher in tall herbaceous (PTG, LEGU, ABP, SHR) and lower in short herbaceous (PSG and PSF).

The results of this study considerably contribute to the sustainable management of grassland resources in the study area. In addition, the results provide a perspective for evaluating current grazing management scenarios and carrying out timely adaptive practices, so as to maintain the ability of the grassland system to perform its ecological functions in the long term.

## Acknowledgments

We thank the staff of Hulunbuir Grassland Ecosystem Research Station for their assistance during field sampling, and our colleagues from the Agricultural Resources and Regional Planning Institute for helping us to make these figures.

## Author Contributions

Conceptualization, Y.M.Z and X.X; Main analysis, Y.M.Z, A.I.A and C.J; Visualization, Y.M.Z and R.Y.; Writing original draft, Y.M.Z and J.S.O; Supervision, X.X and R.Y; Funding and project administration, X.X; Writing: draft editing and Review, X.X and R.Y; Y.M.Z, R.Y, and A.I.A authors have been equally contributed in this paper; all authors have read and agreed to the published version of the manuscript.

